# Excitatory/inhibitory imbalance in autism: the role of glutamate and GABA gene-sets in symptoms and cortical brain structure

**DOI:** 10.1101/2021.12.20.473501

**Authors:** Viola Hollestein, Geert Poelmans, Natalie J Forde, Christian F Beckmann, Christine Ecker, Caroline Mann, Tim Schaefer, Carolin Moessnang, Sarah Baumeister, Tobias Banaschewski, Thomas Bourgeron, Eva Loth, Flavio Dell’Acqua, Declan GM Murphy, Julian Tillmann, Tony Charman, Emily J.H. Jones, Luke Mason, Sara Ambrosino, Rosemary Holt, Sven Bölte, Jan K Buitelaar, Jilly Naaijen

## Abstract

**Background:** The excitatory/inhibitory (E/I) imbalance hypothesis posits that an imbalance between excitatory (glutamatergic) and inhibitory (GABAergic) mechanisms underlies the behavioral characteristics of autism spectrum disorder (autism). However, how E/I imbalance arises and how it may differ across autism symptomatology and brain regions is not well understood.

**Methods:** We used innovative analysis methods - combining competitive gene-set analysis and gene-expression profiles in relation to cortical thickness (CT)- to investigate the relationship between genetic variance, brain structure and autism symptomatology of participants from the EU-AIMS LEAP cohort (autism=360, male/female=259/101; neurotypical control participants=279, male/female=178/101) aged 6 to 30 years. Competitive gene-set analysis investigated associations between glutamatergic and GABAergic signaling pathway gene-sets and clinical measures, and CT. Additionally, we investigated expression profiles of the genes within those sets throughout the brain and how those profiles relate to differences in CT between autistic and neurotypical control participants in the same regions.

**Results:** The glutamate gene-set was associated with all autism symptom severity scores on the Autism Diagnostic Observation Schedule-2 (ADOS-2) and the Autism Diagnostic Interview-Revised (ADI-R) within the autistic group, while the GABA set was associated with sensory processing measures (using the SSP subscales) across all participants. Brain regions with greater gene expression of both glutamate and GABA genes showed greater differences in CT between autistic and neurotypical control participants.

**Conclusions:** Our results suggest crucial roles for glutamate and GABA genes in autism symptomatology as well as CT, where GABA is more strongly associated with sensory processing and glutamate more with autism symptom severity.

## Introduction

Autism Spectrum Disorder (autism) is a neurodevelopmental condition characterized by challenges in social interaction and communication, restricted and repetitive patterns of behavior and/or altered sensory processing (1). Autism is heritable, heterogeneous, and its underlying physiology is still relatively unknown. One influential hypothesis is the excitatory/inhibitory (E/I) imbalance hypothesis, suggesting that imbalance between excitatory (predominantly glutamatergic) and inhibitory (predominantly GABAergic (γ-aminobutyric acid)) mechanisms in the brain underlies autism symptomatology (2). Causal links for this have been suggested, and both overexcitation and overinhibition have been associated with autism before (2–6). However, understanding the mechanisms of *how* E/I imbalance is underlying autism symptomatology is complex. The heterogeneity and polygenic nature of autism, and previous opposing findings of E/I imbalance, may be evidence of differential involvement across autism characteristics or brain regions.

Mechanisms of E/I imbalance may have genetic underpinnings. Autism is a polygenic condition where several genetic variants together give rise to the expression of the phenotype, and progress in identifying common genetic variants associated with autism includes genes encoding for proteins involved in glutamate and GABA receptors and transporters (7–10). Several studies have suggested glutamatergic and GABAergic genetic links to behavioral autism phenotypes (3,4,11–13). These phenotypes have been linked to changes in glutamate and GABA concentrations in the brain as well (4,14).

Anatomical differences may also be affected by E/I imbalance, as glutamate and GABA receptors play a role in dendritic growth, a process with genetic underpinnings found to be altered in autism (15–17). Dendrite growth is also linked to cortical thickness (CT) (18,19). Altered dendrite growth, and associated genes, have been linked to autism symptomatology, especially repetitive behaviors (15,20,21). Increased and decreased CT has also been found in autism, mainly in fronto-temporal, fronto-parietal, limbic areas and fronto-striatal circuits (22–28). Further, cell-type specific gene-expression has been shown to be associated with CT (18,29). This suggests that differences in CT may be partly due to genetic underpinnings of the glutamate and GABA systems, which could underlie autism symptomatologies. However, this has not yet been investigated in genes involved in regulating excitatory and inhibitory signaling.

To deepen the understanding of relationships between these measures, this study uses a hypothesis driven approach within the E/I imbalance framework. We use a competitive gene-set analysis approach (30–32) investigating the role of glutamatergic and GABAergic genes in behavioral autism phenotypes and CT. This method tests whether the genes in a gene-set (here glutamate and GABA gene-sets) are more strongly correlated with a phenotype than the other genes in the genome. This method considers several genetic variants in the same analysis, which increases both the power of the study and the phenotypic variance explained compared to methods investigating genetic variants separately. This method has shown utility in other neurodevelopmental disorders showing aggregated genetic effects rather than single candidate-gene associations (20,33,34). To further examine the role of glutamatergic and GABAergic genes affecting differences in brain structure, we used a gene-expression profile approach (29,35) to investigate the association between gene-expression of glutamatergic and GABAergic genes and differences in CT between autistics and neurotypical controls (NTC) throughout the cortex. With gene-expression profile analysis we investigate whether altered expression of the glutamate and GABA genes are correlated with differences in CT profiles between autism and NTC across the brain, further strengthening associations the gene-set analysis may give.

By combining these approaches, we can deepen our understanding of the links between glutamate and GABA pathway signaling genes, differences in CT and behavioral autism characteristics. From the gene-set analyses we expect a differential involvement of glutamatergic and GABAergic gene-sets in behavioral autism phenotypes and brain structure. The association with gene-expression of those gene-sets is exploratory to investigate whether differences in CT is associated with expression of glutamatergic and GABAergic genes across brain regions (29,35).

## Methods and Materials

### Participants

Autistic (n=360) and neurotypical control participants (NTC; n=279) were part of the Longitudinal European Autism Project (LEAP) within the EU-AIMS clinical research programme (https://www.aims-2-trials.eu/) (36–38). Phenotypic, genetic and brain imaging data were collected at six study centers across Europe: Institute of Psychiatry, Psychology and Neuroscience, King’s College London (IoPPN/KCL, UK), Autism Research Centre, University of Cambridge (UCAM, UK), University Medical Centre Utrecht (UMCU, Netherlands), Radboud University Nijmegen Medical Centre (RUNMC, Netherlands), Central Institute of Mental Health (CIMH, Germany), and the University Campus Bio-Medico (UCBM) in Rome, Italy.

Inclusion criteria for the autism group were an existing diagnosis of autism and an age-range between 6 to 30 years. Symptoms were additionally assessed using the Autism Diagnostic Observation Schedule (ADOS-2; (39)) and the Autism Diagnostic Interview-Revised (ADI-R; (40)). For the NTC, exclusion criterion comprised parent-or self-report of any psychiatric disorder. Individuals who had a normative T-score of 70 or higher on the Social Responsiveness Scale Second Edition (SRS-2) were excluded. Some individuals in the autism and NTC groups had mild intellectual disability (ID) (autism=53, NTC=25), defined as an IQ score between 40 and 74. For further details of the recruitment of participants in this study see (24,36,37).

### Phenotypic measures

The phenotypic measures used were part of a larger test battery (see (37)). Here we included three questionnaires regarding the main autism symptomatology; the Social Responsiveness Scale-Revised (SRS-2) (41), the Repetitive Behavior Scale-Revised (RBS-R) (42), and the Short Sensory Profile (SSP) (43). For these questionnaires, we used self-or parent-report ratings, depending on age and diagnostic group.

### Genotyping

Genotyping was performed at the Centre National de Recherche en Génomique Humaine (CNRGH) using the Infinium OmniExpress-24v1 BeadChip Illumina. Sample quality controls such as sex check (based on the X chromosome homozygosity rate or the median of the Log R ratio of the X and Y chromosomes), Mendelian errors (transmission errors within full trios) and Identity By State were performed using PLINK 1.90. Imputation of 17 million SNPs was performed using the 700k genotyped SNPs on the Michigan Imputation Server (44). The HRC r1.1 2016 reference panel for a European population was used, as the majority of individuals in the LEAP cohort were from European ancestry. Only autosomes were imputed. Linkage disequilibrium-based SNP pruning was done for SNPs with a MAF>1% and SNPs with an R2 < 0.1 in windows of 500kb were selected. This resulted in 547 participants with genotypic data.

Two types of gene-sets were used both for glutamate and GABA, which resulted in a total of four gene-sets being tested on each phenotype of interest. The first and largest gene-set type consists of genes encoding proteins involved in the glutamatergic and GABAergic signaling pathways, these will be referred to as the glu-pathway (n_genes_=72) and GABA-pathway (n_genes_=124) gene-sets. The selection of the genes in these sets was based on Ingenuity Pathway Analysis software (http://www.ingenuity.com), a frequently updated database for genetic pathway analysis. Supplemental tables S1 and S2 show an overview of the included genes. Additionally, we investigated associations with genes encoding glutamate/GABA receptors and transporters specifically because of their more direct role in neurotransmitter signaling (45), referred to as glu-RT (n=32) and GABA-RT (n=26).

### Neuroimaging data

Structural brain images were acquired on 3 T MRI scanners at all sites, with T1-weighted MPRAGE sequence (TR=2300ms, TE=2.93ms, T1=900ms, voxels size=1.1×1.1×1.2mm, flip angle=9°, matrix size=256×256, FOV=270mm, 176 slices). For a summary of scanner details and acquisition parameters at each site, see Table S3 in the supplemental information.

For each image a model of the cortical surface was computed using FreeSurfer v6.0 (https://surfer.nmr.mgh.harvard.edu/), using a fully automated and validated procedure (46–49). Subsequently, each reconstructed surface went through strict quality assessments, described in detail in (24). This resulted in parcellated regional CT measures for all the 639 participants included in our study, with 34 regions in each hemisphere using the Desikan-Killiany atlas (50).

### Gene-expression data

Gene-expression data were acquired from the Allen Human Brain Atlas (AHBA) (51). These data were obtained from six post-mortem donors (five males, one female), where the brains of the donors were sampled to anatomical locations in MNI152 space. These whole-brain gene-expression data are open source and downloadable from the Allen Institute for Brain Science; http://www.brain-map.org). For more details on how these data were obtained, see (51).

In previous work, these AHBA gene-expression data were aggregated across donors and mapped onto the 68 FreeSurfer areas, 34 per hemisphere (35,50). This was done by taking the gene-expression data from AHBA, containing all voxels of gene-expression values across donors and mapping these voxels onto the FreeSurfer areas. All voxels in each area were then median-aggregated across donors. As data was available for all AHBA donors in the left hemisphere (and only from two donors in the right hemisphere), only left hemisphere data was included in our analysis.

These gene-expression profiles were then used in the two-step procedure described by (35) to select the most consistent profiles for inclusion in our analyses. First, the correlations of gene-expressions to the median expression values across donors were calculated, and the genes showing consistent correlation profiles were selected (donor-to-median correlation rho >0.446). Secondly, we used data from the BrainSpan Atlas, where gene-expression data in a wide age-range of donors are available (www.brainspan.org). Donors were selected within the age range of our LEAP dataset (6-30 years), which gave us 9 donors (male/female = 5/4). We calculated correlations to the median expression values in the 11 cortical regions in the AHBA-to-FreeSurfer data that were also included in the BrainSpan Atlas, using scripts by (29). We then selected genes that correlated between the profiles of the two atlases higher than r=0.52 (one-sided *test p* < 0.05), which resulted in 2293 genes available in total. The overlap with the gene-sets left 23 genes in our glutamate pathway gene-set, and 39 genes in our GABA pathway gene-set. The number of overlapping genes with our RT gene-sets were too few and were therefore not included. The median expression profiles across regions for these genes constitute the interregional gene-expression profiles used in our analyses.

### Analyses

All analyses included age, sex, IQ and site as covariates. All tests were corrected using false discovery rate (FDR) unless otherwise described, therefore the *p*-value significance threshold was set to *p* < 0.05.

### Competitive gene-set analysis

To investigate associations of the glutamate and GABA (pathway, and RT) gene-sets with the phenotypes of interest (SRS-2 total score, RBS-R total score, SSP total score, and ADOS-2 and ADI-R (the last two for the autism group only)), we performed competitive gene-set analysis using MAGMA (Multi-marker Analysis of GenoMic Annotation) software (version 1.07b, (30)). We also investigated glutamatergic and GABAergic gene-set associations with regional CT.

The analysis was performed in two steps. First, gene-based *p*-values were calculated for each gene (excluding genes located on the X-chromosome, see supplemental tables S1 and S2) on our phenotypes of interest, using a multiple linear principal components regression using F-tests. Second, we tested the association of the set, aggregating the gene-based *p*-values using competitive analysis. This gene-set analysis is done with an intercept-only linear regression model for the gene-set, which tests whether the genes in a gene-set are more strongly correlated with the phenotype of interest than all other genes in the genome (30).

### Cortical thickness and clinical phenotypes

To test associations between CT and our phenotypes of interest (continuous measures of core autism-symptomatology), we used linear regression models in the R-software package (52). This was done in the left hemisphere only, due to the expression profile analysis being performed only in the left hemisphere. Sex, linear and quadratic effects of age, full scale IQ, site and mean CT were used as covariates in all our models as described in (24).

### Expression profiles

To investigate correlations between glutamatergic and GABAergic gene-expression and brain structure, we used the gene-expression profiles (29,35) approach. This uses correlation across interregional profiles of CT with interregional profiles of gene-expression (18,29,53). The interregional profiles of MRI-derived CT-differences were created by calculating the average CT in the autism group, across regions from our LEAP sample, and subtracting it from the average CT of the NTC (matched for age, sex and IQ to make the sample as homogeneous as possible, n=279 in both groups). This gave us one average value of CT-difference per region, constituting the interregional profiles of CT. Only left hemisphere data of both gene-expression and CT-difference was used in in subsequent analyses.

Interregional profiles of gene-expressions for our pathway gene-sets were calculated by taking the median of the donor expressions, described earlier, for each gene in our gene-sets per region. As the filtered (see above) AHBA data did not have expression data available for all genes in our gene-sets, the number of genes in the gene-sets were smaller compared to previous analyses (glu-pathway=23, GABA-pathway=39). The interregional profiles of the genes in our glutamate and GABA pathway gene-sets were then correlated with the CT-difference interregional profiles, which provided a distribution of correlation coefficients per gene-set. The distributions of correlation coefficients between the gene-expression and CT-difference interregional profiles were then tested for significance using a resampling approach of 10,000 random samples, as described in (18,29). In this approach, a random set of genes of the same size as the set being tested, was selected (from the 2293 available) 10,000 times with the average correlation each time being used to create a null distribution. A two-tailed significance test was used to test the gene-set of interest against the null distribution.

As CT is known to be strongly associated with age (54), we repeated these analyses in separate age-groups of our sample (children: 6-11 years, adolescents: 12-17 years, adults: 18 years and older). We also replicated this gene-expression analysis in an independent dataset, using structural imaging data of CT from the multi-site open-source ABIDE database (55). We included participants in the same age range as our own sample (6-30 years), which resulted in data from 874 participants matched for age, sex and IQ (autism =437, male/female = 385/52, NTC =437, male/female = 349/88). Details on these results can be found in the supplemental information.

## Results

### Demographics

Demographic and clinical characteristics are shown in Table 1. No differences were found between the autism and NTC group in age. The autism group had a higher female-to-male ratio compared to NTC and NTC had a significantly higher IQ than the autism group. As expected, the autism group had significantly higher scores on the SRS-2 and RBS-R and scored lower in the SSP (where lower scores indicate higher sensory sensitivity). Information on medication use can be found in Table S4 in the supplemental information.

**Table 1.**
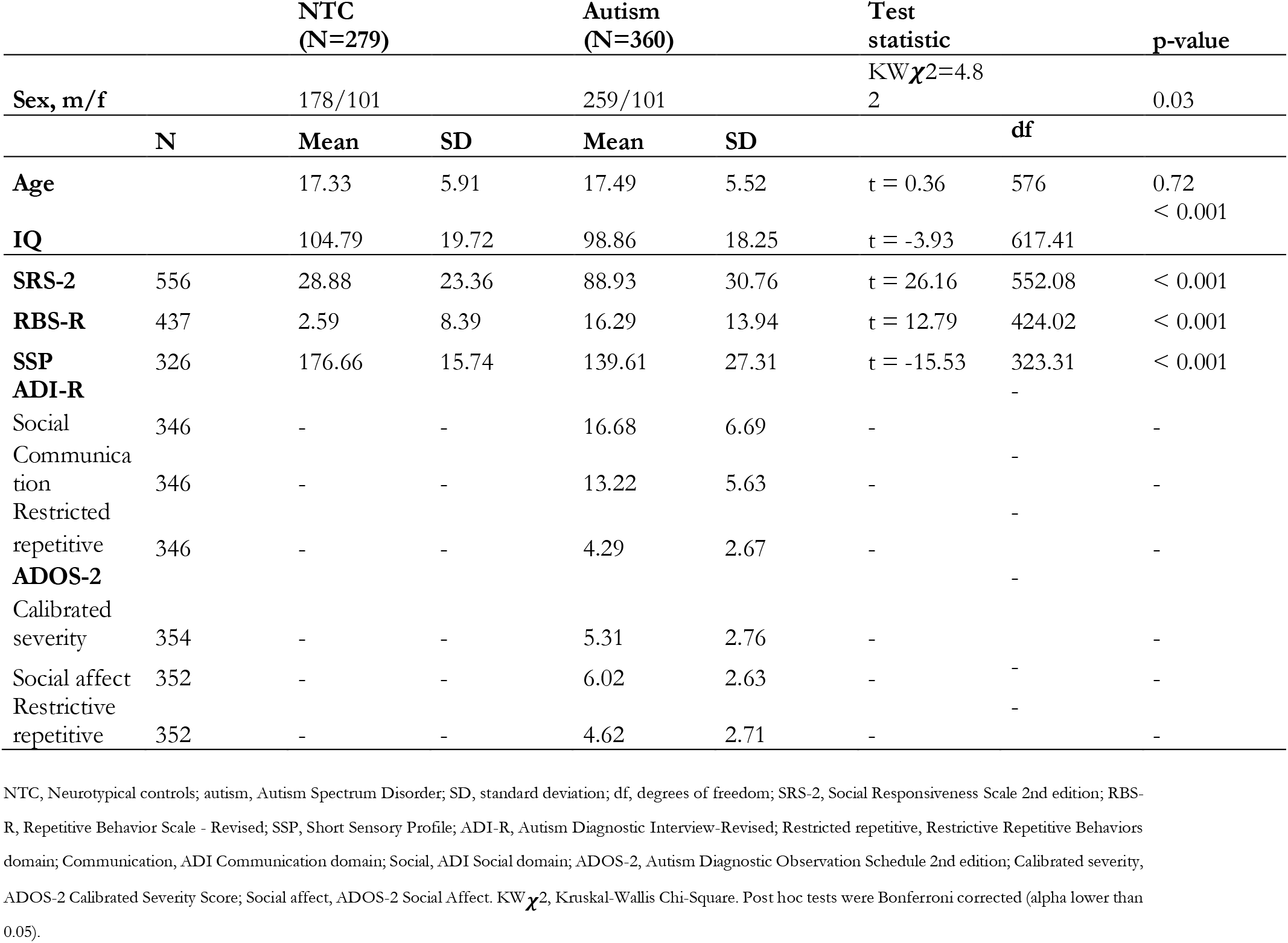
Demographic and clinical characteristics

### Competitive gene-set analysis

Significant associations between the glu-pathway gene-set (N=72 genes) and all the ADI-R and ADOS-2 subscales (all *p*_*FDR*_-values < 0.05) were found. No significant associations were found for SRS-2, RBS-R or SSP. The GABA-pathway and GABA-RT gene-sets were nominally significantly associated with SSP total scores (GABA-pathway *p*_FDR_ = 0.07, GABA-RT *p*_FDR_=0.06). However, this did not survive FDR-correction, see Table 2 for more details. To investigate whether the trend association of GABA and SSP could be explained by variance within the SSP, we performed post-hoc analyses on all SSP subscales, which were FDR corrected for n=7 subscales. Significant associations of both the GABA-pathway and GABA-RT sets on all SSP subscales were found (no significance with the glutamate gene-sets), which all survived FDR-correction (*q*-values < 0.05), and had similar regression coefficients (β). For more details see Table S5 in the supplemental information.

**Table 2.** Glutamate and GABA and phenotypes competitive gene-set analysis results

| **Glutamate: Pathway gene-set (N=72)** | **BETA** | **P** | **P<sub>FDR</sub>** | **SE** |
| --- | --- | --- | --- | --- |
| SRS | 0.076 | 0.244 | 0.244 | 0.109 |
| RBS-R | 0.112 | 0.141 | 0.221 | 0.104 |
| SSP | 0.106 | 0.148 | 0.221 | 0.101 |
| Diagnosis | 0.190 | 0.033 | - | 0.103 |
| **ADI communication domain** | **0.213** | **0.019** | **0.038** | **0.103** |
| **ADI restricted and repetitive behaviors domain** | **0.213** | **0.019** | **0.019** | **0.103** |
| **ADI social domain** | **0.213** | **0.019** | **0.019** | **0.103** |
| **ADOS restricted and repetitive behaviors** | **0.242** | **0.009** | **0.019** | **0.103** |
| **ADOS social affect** | **0.242** | **0.009** | **0.019** | **0.103** |
| **ADOS total score** | **0.242** | **0.009** | **0.019** | **0.103** |
| **Glutamate: Receptors/transporters gene-set (N=31)** |  |  |  |  |
| SRS | -0.010 | 0.525 | 0.539 | 0.170 |
| RBS-R | -0.016 | 0.539 | 0.539 | 0.163 |
| SSP | 0.184 | 0.122 | 0.364 | 0.158 |
| Diagnosis | 0.099 | 0.270 | - | 0.160 |
| ADI communication domain | 0.104 | 0.257 | 0.323 | 0.160 |
| ADI restricted and repetitive behaviors domain | 0.104 | 0.258 | 0.323 | 0.160 |
| ADI social domain | 0.104 | 0.257 | 0.323 | 0.160 |
| ADOS restricted and repetitive behaviors | 0.211 | 0.094 | 0.323 | 0.160 |
| ADOS social affect | 0.212 | 0.094 | 0.323 | 0.160 |
| ADOS total score | 0.211 | 0.094 | 0.323 | 0.160 |
| **GABA: Pathway gene-set (N=124)** |  |  |  |  |
| SRS | -0.052 | 0.728 | 0.728 | 0.085 |
| RBS-R | -0.019 | 0.591 | 0.728 | 0.082 |
| **SSP** | **0.157** | **0.024** | 0.071 | 0.079 |
| Diagnosis | 0.046 | 0.285 | - | 0.080 |
| ADI communication domain | 0.054 | 0.251 | 0.291 | 0.080 |
| ADI restricted and repetitive behaviors domain | 0.054 | 0.251 | 0.291 | 0.080 |
| ADI social domain | 0.054 | 0.251 | 0.291 | 0.080 |
| ADOS restricted and repetitive behaviors | 0.044 | 0.291 | 0.291 | 0.080 |
| ADOS social affect | 0.044 | 0.291 | 0.291 | 0.080 |
| ADOS total score | 0.044 | 0.290 | 0.291 | 0.080 |
| **GABA: Receptors/transporters gene-set (N=23)** |  |  |  |  |
| SRS | 0.094 | 0.331 | 0.497 | 0.215 |
| RBS-R | -0.328 | 0.945 | 0.945 | 0.205 |
| **SSP** | **0.404** | **0.021** | 0.063 | 0.199 |
| Diagnosis | 0.151 | 0.227 | - | 0.202 |
| ADI communication domain | 0.141 | 0.243 | 0.268 | 0.202 |
| ADI restricted and repetitive behaviors domain | 0.140 | 0.244 | 0.268 | 0.202 |
| ADI social domain | 0.141 | 0.243 | 0.268 | 0.207 |
| ADOS restricted and repetitive behaviors | 0.125 | 0.268 | 0.268 | 0.202 |
| ADOS social affect | 0.125 | 0.267 | 0.268 | 0.202 |
| ADOS total score | 0.126 | 0.267 | 0.268 | 0.202 |

N, number of genes in analysis. Diagnosis was indicated as a binary variable. SRS-2, Social Responsiveness Scale, Second Edition; ; RBS-R, Repetitive Behavior Scale-Revised; SSP, Short Sensory Profile; ADI, Autism Diagnostic Interview-Revised; ADOS, Autism Diagnostic Observation Schedule; P<sub>FDR</sub> p-value corrected using False discovery rate (FDR); SE, standard error of the regression coefficient. Significant results (p<sub>FDR</sub> < 0.05) marked in bold.

No other significant associations were found with the other phenotypes of interest or with the glu-RT nor GABA-RT gene-sets. As the associations with ADI-R and ADOS-2 were only available in the autism group, we also performed competitive gene-set analysis with the SRS-2, RBS-R and SSP in the autism group separately, investigating whether there were additional effects only seen in this group. This gave no significant associations. We additionally investigated gene-set association with CT in the FreeSurfer cortical regions in the left hemisphere. There were some nominally significant (nominal *p*-values <0.05) associations, although none survived FDR-correction. The details of these results can be seen in supplemental tables S6-S9.

### Cortical thickness and phenotypes

Vertex-wise comparisons in CT between autism participants and NTC has been performed previously on this cohort, described in a recent manuscript (24). Here we focused on associations with continuous measures related to autism symptomatology using parcellated regional measures, by using SRS-2, RBS-R and SSP questionnaire scores across the entire sample and sub-scales of ADI-R and ADOS-2 in the autism group. We found a positive association between CT in the frontal pole and restricted and repetitive behaviors as reflected by the RBS-R total score (b = 0.05, t = 3.33, *p*_FDR_ = 0.03). To investigate whether this was driven by the autism or NTC group, we performed the analysis separately for these groups, which showed that this effect was driven by the autism group. There were also nominal negative associations between ADI-R subscales and CT in precuneus, and positive associations between the total ADOS-2 score and social domain subscale and CT in the insula. However, these did not survive FDR-correction.

### Gene-expression profiles

The interregional profiles of group differences in CT (autism-NTC) was associated with the interregional profile of gene expression for both glutamate (N=23) and GABA-pathway (N=39) gene-sets (glu-pathway: t=-2.58, *p*_FDR_ = 0.020, d=0.97, GABA-pathway: t=-2.42, *p*_FDR_ = 0.020, d=0.76). As shown in Figure 1, regions with greater gene expression of both glutamatergic and GABAergic genes showed greater differences in CT between autistic and neurotypical participants. Repeating these analyses in the different age groups showed that this effect was mainly present in the adult group. Additionally, the ABIDE replication analyses showed some similar but not a complete replication of our results. Details can be found in the supplemental information and Figure S4.

**Figure 1:**
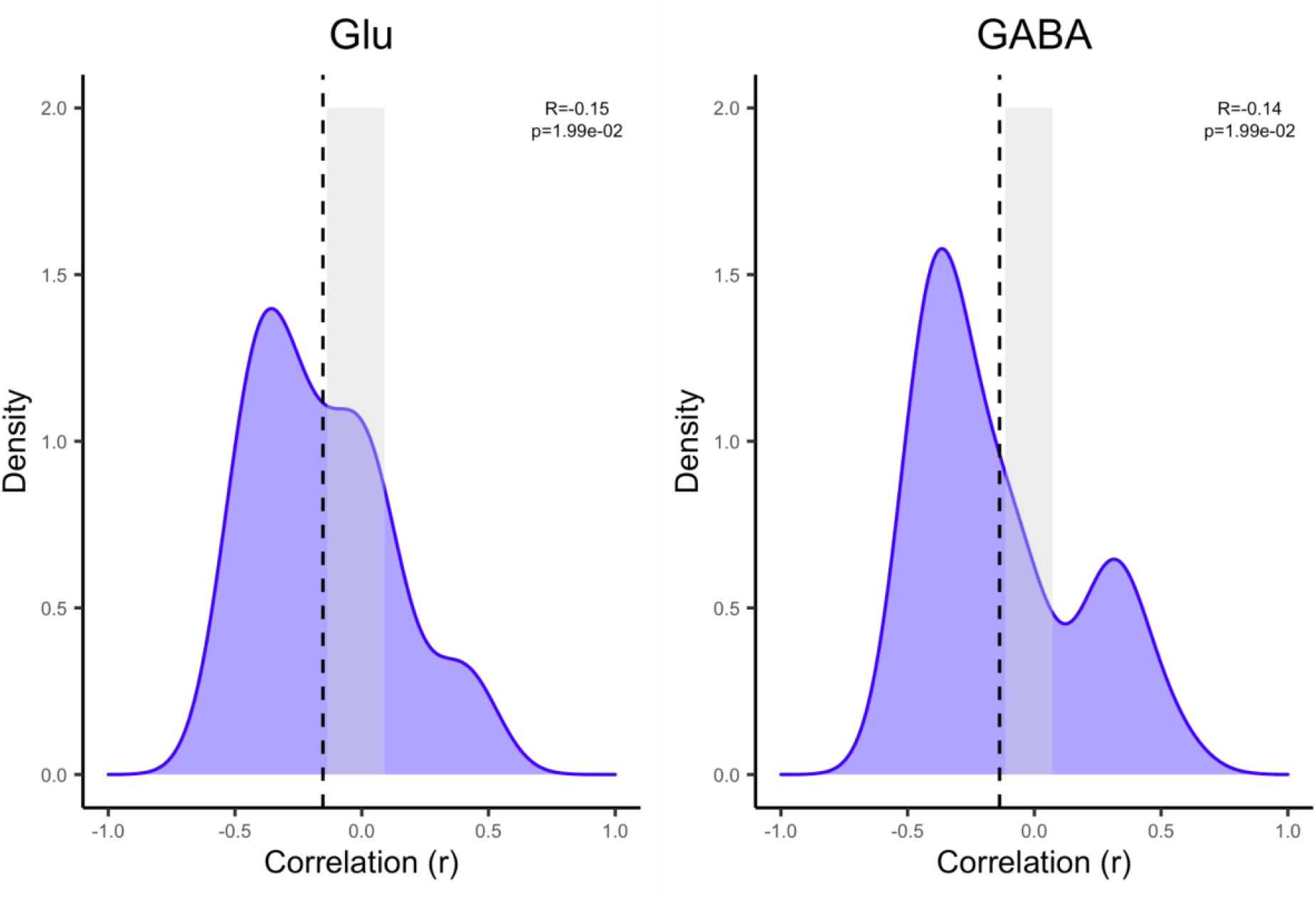
Distributions of correlation coefficients between cortical thickness and gene-expression Distributions of the inter-regional correlation coefficients between differences in cortical thickness (CT) and profiles of gene-expression. The CT-difference profile was obtained from our LEAP data, and the expression profiles from the Allen Human Brian Atlas (AHBA), in our glutamate-pathway and GABA-pathway gene-sets. The x-axes show the correlation coefficient between CT-difference and expression profile for the gene-set; the y-axes show the estimated probability density for the correlation coefficients; the vertical dashed-lines indicates the average expression-CT difference correlation coefficient across all the marker genes in a gene-set; and the edges of the gray boxes indicates the 2.5% and 97.5%-critical values obtained from the empirical null distribution of the average expression-thickness correlation coefficient. If a vertical line sits outside the gray box, it implies that there is a significant association between gene-set and differences in CT at the unadjusted 5% significance level.

**Figure 2:**
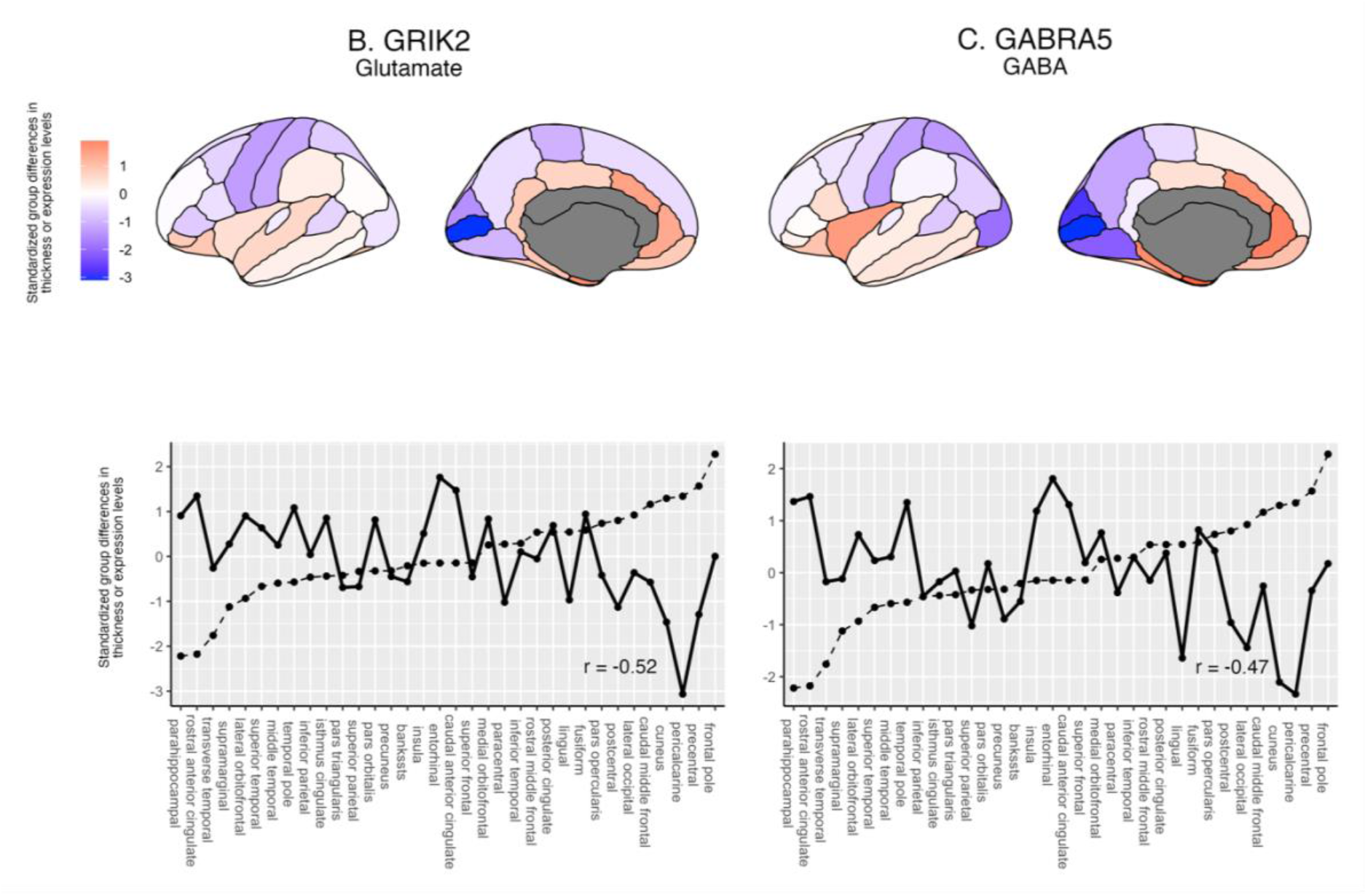
Gene expression and cortical thickness from highest correlating genes Lateral and medial views of differences in cortical thickness (CT) between autistic and neurotypical control participants, and gene-expression levels of the genes from the glutamate and GABA (pathway) gene-sets with highest (negative) correlation. Plots on the bottom row show standardized profiles of CT-differences between autism and neurotypical controls (dotted lines) and the gene-expression (solid lines) for the respective genes of the gene-sets. FreeSurfer regions on the x-axis are ordered from low to high thickness. Figures were created using ggplot2 (Ginestet, 2011) and ggseg (Mowinckel & Vidal-Piñeiro, 2020).

## Discussion

This is the first study to investigate the role of glutamate and GABA genes in behavioral phenotypes and brain structure in autism. The most important takeaway of our results is that glutamate and GABA gene-sets were differently associated with autism symptoms, with glu-pathway genes being associated with autism symptom severity on the ADI-R and ADOS-2, and GABA genes showing associations with sensory processing subscales of the SSP. Additionally, regions with greater gene expression of glutamatergic and GABAergic genes showed greater differences in CT between autism and controls. These results provide a better understanding of the mechanistic underpinnings of the E/I imbalance hypothesis of autism, by supporting the notion that E/I imbalance varies across behavioral autism characteristics and differences in CT between autism and NTC groups.

The findings of associations between the glu-pathway gene-set with ADI-R and ADOS-2 subscale scores, and the trend associations of ADI-R and ADOS-2 scores with CT in the precuneus and insula, areas known to be involved in somatosensory and visuospatial processing, interoception and self-reflection (56,57), suggest that glutamate genes may be linked to broader autism characteristics. The regression coefficient (β) of the associations range between 0.02 and 0.04 (see Table 2), similarly across both ADI-R and ADOS-2, pointing towards more general associations with autism. Additionally, the associations of GABA-pathway and GABA-RT gene-sets on all SSP subscales indicate that GABA genes are linked to sensory processing alterations. These findings are consistent with previous results, as altered GABA concentrations in autism have been associated with altered sensory processing previously (58–60).

The lack of significant associations between the glutamate and GABA gene-sets with repetitive behaviors and social responsiveness (RBS-R and SRS-2) may be considered surprising, as previous studies have found links between glutamate and GABA concentrations in several brain regions and/or metabolite altering drugs, with social responsiveness and repetitive behaviors (61– 66). However, studies investigating in vivo measures of alterations of glutamate and GABA in autism have had inconsistent results that could be due to several factors; the heterogeneity of autism, differences in study populations and brain regions investigated, or differences in processing pipelines during analysis. Furthermore, here we focused on behavioral autism characteristics and genetic information, not in vivo brain concentrations of glutamate and GABA.

Regarding the CT associations with autism characteristics, increased repetitive behaviors measured across the whole group with the RBS-R, were associated with increased CT in the frontal pole. Measures from the ADI-R and ADOS-2 diagnostic tools (in the autism group only) were differently associated with CT in the precuneus and insula, although this was only at trend-level. We did not find associations of CT with SSP scores, although previous work on this dataset did find differences in CT between subgroups of autism (24), likely due to the subgrouping of participants that was not done here. Additionally, interregional variation in expression of glutamatergic and GABAergic genes was associated with the group differences in CT; regions with greater expression of both glutamate and GABA genes showed greater differences in CT between autism and NTC. Separating by age groups, these associations were mainly replicated in adults, and was in the opposite direction in adolescents. This suggests there may be important differences in trajectories across development, which needs to be investigated further. More studies including those on metabolite concentrations are needed to draw further conclusions about these relationships.

Our results need to be interpreted with caution, as the presence of glutamate and GABA protein encoding genes does not directly translate to metabolite concentrations, and genetic alterations might not translate to a common phenotype across individuals (11). Additionally, genes differ in coding for loss-or gain-of function, leading to reduced or increased protein function, further complicating any interpretation of *direction* of glutamate and GABA involvement in autism symptoms. However, our results strongly indicate critical roles of glutamate and GABA genes in these specific phenotypes and that the link between these measures needs to be investigated in more detail to increase our understanding of the mechanisms connecting genetics, glutamate and GABA neurotransmitters and autism symptomatology. More direct investigations of the E/I imbalance hypothesis are needed in order to investigate excitation and inhibition in vivo in relation to brain functioning. Promising new techniques combining different imaging methods, causal discovery analysis, and pharmacological interventions and longitudinal studies, will allow us to do this in the future through which we hope to further increase our understanding of how brain chemical imbalance is associated with functioning. Ultimately, E/I balance may be manipulated using glutamate- and/or GABA-influencing pharmacological treatments. One study already showed decreased glutamate and GABA concentrations after bumetanide treatment to be positively associated with autism symptom improvement (67).

Strengths of this study were the combination of genetic, structural and phenotypic data from the same cohort, which gave us the opportunity to for the first time analyze these data together. Another strength was the relatively large number of participants available giving us more confidence in our results. There were also some limitations. Firstly, there were fewer females than males included in this study, a common problem in autism research. Furthermore, the gene-expression data were only used in the left hemisphere. However, the gene-expression data used in the expression profile analyses was robust and only included if the interregional profiles were similar across another dataset (BrainSpan), increasing the confidence in the robustness of these profiles. Another limitation is that the AHBA donors were all neurotypical controls, and we do not know whether genes are expressed differently in autism. We also did not fully replicate our gene-expression profile results in the independent ABIDE data set (see the supplemental information and Figure S4). However, in the GABA-pathway gene-set there was a similar direction of possible effects, although not significant. This shows that heterogeneity of the autism sample and neurotypical controls have a large influence on the results.

In conclusion, we found that GABA genes are associated with sensory processing while glutamate genes are associated with behavioral autism characteristics more broadly, and that increased expression of these glutamate and GABA genes are associated with larger differences in CT between autistics and NTC. This support the hypothesis that the influence of E/I imbalance varies across autism phenotypes and brain regions, suggesting that glutamate and GABA genes play different roles underlying different autism phenotypes. We also showed the importance of linking structural brain measures, genetic and behavioral phenotype data together to gain a deeper understanding of possible E/I imbalance mechanisms in autism.

## Acknowledgements

This work has been supported by the EU-AIMS (European Autism Interventions) and AIMS-2-TRIALS programmes which receive support from Innovative Medicines Initiative Joint Undertaking Grant No. 115300 and 777394, the resources of which are composed of financial contributions from the European Union’s FP7 and Horizon2020 Programmes, and from the European Federation of Pharmaceutical Industries and Associations (EFPIA) companies’ in-kind contributions, and AUTISM SPEAKS, Autistica and SFARI; by the Horizon2020 supported programme CANDY Grant No. 847818) and a Veni grant from the Netherlands organization for scientific research (NWO) under grant number VI.Veni.194.032 awarded to J Naaijen.

The funders had no role in the design of the study; in the collection, analyses, or interpretation of data; in the writing of the manuscript, or in the decision to publish the results. Any views expressed are those of the authors and not necessarily those of the funders.

