## Supplemental information for "Excitatory/inhibitory imbalance in autism: the role of glutamate and GABA gene-sets in symptoms and cortical brain structure"

**Hollestein et al.**

**Table S1:** Summary table of all genes in glutamate gene-set included in analyses.

| Gene name | Entrez gene ID | Chromosome | Start position | End position | strand | NSNPS |
| --- | --- | --- | --- | --- | --- | --- |
| ABAT | 18 | 16 | 8768444 | 8878432 | + | 1013 |
| ALDH5A1 | 7915 | 6 | 24495197 | 24537435 | + | 297 |
| CALM1 | 801 | 14 | 90863327 | 90874619 | + | 47 |
| CALML5 | 51806 | 10 | 5540658 | 5541533 | - | 6 |
| CAMK4 | 814 | 5 | 110559947 | 110830584 | + | 1538 |
| DLG4 | 1742 | 17 | 7093209 | 7123369 | - | 102 |
| GAD1 | 2571 | 2 | 171673200 | 171717661 | + | 172 |
| GAD2 | 2572 | 10 | 26505236 | 26593491 | + | 579 |
| GLS | 2744 | 2 | 191745547 | 191830278 | + | 290 |
| GLUD1 | 2746 | 10 | 88809959 | 88854776 | - | 186 |
| GLUD2 | 2747 | X | 120181462 | 120183796 | + |  |
| GLUL | 2752 | 1 | 182350839 | 182361341 | - | 55 |
| GNB1 | 2782 | 1 | 1716725 | 1822552 | - | 250 |
| GNB1L | 54584 | 22 | 19775932 | 19842462 | - | 369 |
| GNB2 | 2783 | 7 | 100271363 | 100276792 | + | 19 |
| GNB3 | 2784 | 12 | 6949375 | 6956564 | + | 34 |
| GNB5 | 10681 | 15 | 52413123 | 52483565 | - | 486 |
| GNG10 | 2790 | 9 | 114423851 | 114432526 | + | 50 |
| GNG11 | 2791 | 7 | 93551016 | 93555826 | + | 32 |
| GNG12 | 55970 | 1 | 68167149 | 68299436 | - | 702 |
| GNG13 | 51764 | 16 | 848041 | 850733 | - | 33 |
| GNG2 | 54331 | 14 | 52327022 | 52436518 | + | 794 |
| GNG3 | 2785 | 11 | 62475066 | 62476678 | + | 5 |
| GNG4 | 2786 | 1 | 235710985 | 235814054 | - | 543 |
| GNG5 | 2787 | 1 | 84964006 | 84972262 | - | 37 |
| GNG7 | 2788 | 19 | 2511218 | 2702746 | - | 1041 |
| GOT1 | 2805 | 10 | 101156627 | 101190530 | - | 146 |
| GOT1L1 | 137362 | 8 | 37791799 | 37797664 | - | 17 |
| GOT2 | 2806 | 16 | 58741035 | 58768246 | - | 229 |
| **GRIA1** | **2890** | **5** | **152870084** | **153193429** | **+** | 1819 |
| **GRIA2** | **2891** | **4** | **158141736** | **158287227** | **+** | 425 |
| **GRIA3** | **2892** | **X** | **122317996** | **122624766** | **+** |  |
| **GRIA4** | **2893** | **11** | **105480800** | **105852819** | **+** | 1505 |
| GRID1 | 2894 | 10 | 87359312 | 88126250 | - | 4622 |
| GRID2 | 2895 | 4 | 93225453 | 94695707 | + | 7119 |
| **GRIK1** | **2897** | **21** | **30909254** | **31312282** | **-** | 2258 |
| **GRIK2** | **2898** | **6** | **101841584** | **102517958** | **+** | 3720 |
| **GRIK3** | **2899** | **1** | **37261128** | **37499844** | **-** | 963 |
| **GRIK4** | **2900** | **11** | **120382465** | **120859514** | **+** | 2775 |
| **GRIK5** | **2901** | **19** | **42502468** | **42574278** | **-** | 138 |
| **GRIN1** | **2902** | **9** | **140033609** | **140063214** | **+** | 86 |
| **GRIN2A** | **2903** | **16** | **9847265** | **10276611** | **-** | 3419 |
| **GRIN2B** | **2904** | **12** | **13713684** | **14133022** | **-** | 2569 |
| **GRIN2C** | **2905** | **17** | **72838162** | **72856966** | **-** | 93 |
| **GRIN2D** | **2906** | **19** | **48898132** | **48948188** | **+** | 222 |
| **GRIN3A** | **116443** | **9** | **104331634** | **104500862** | **-** | 942 |
| **GRIN3B** | **116444** | **19** | **1000437** | **1009723** | **+** | 108 |
| GRINA | 2907 | 8 | 145064226 | 145067596 | + | 9 |
| GRIP1 | 23426 | 12 | 66741178 | 67463014 | - | 4124 |
| **GRM1** | **2911** | **6** | **146286032** | **146758782** | **+** | 2121 |
| **GRM2** | **2912** | **3** | **51741081** | **51752629** | **+** | 16 |
| **GRM3** | **2913** | **7** | **86273230** | **86494193** | **+** | 1110 |
| **GRM4** | **2914** | **6** | **33989623** | **34123399** | **-** | 1020 |
| **GRM5** | **2915** | **11** | **88237256** | **88796846** | **-** | 3817 |
| **GRM6** | **2916** | **5** | **178405328** | **178422124** | **-** | 141 |
| **GRM7** | **2917** | **3** | **6902802** | **7783218** | **+** | 5656 |
| **GRM8** | **2918** | **7** | **126078652** | **126892428** | **-** | 4521 |
| HOMER1 | 9456 | 5 | 78669647 | 78809659 | - | 705 |
| HOMER2 | 9455 | 15 | 83517729 | 83654905 | - | 736 |
| HOMER3 | 9454 | 19 | 19040010 | 19052041 | - | 42 |
| PICK1 | 9463 | 22 | 38453262 | 38471708 | + | 92 |
| SLC17A1 | 6568 | 6 | 25783125 | 25832287 | - | 297 |
| SLC17A2 | 10246 | 6 | 25912982 | 25930954 | - | 109 |
| **SLC17A6** | **57084** | **11** | **22359667** | **22401049** | **+** | 208 |
| **SLC17A7** | **57030** | **19** | **49932655** | **49945617** | **-** | 39 |
| **SLC17A8** | **246213** | **12** | **100750857** | **100815837** | **+** | 347 |
| **SLC1A1** | **6505** | **9** | **4490427** | **4587469** | **+** | 544 |
| **SLC1A2** | **6506** | **11** | **35272752** | **35441610** | **-** | 1155 |
| **SLC1A3** | **6507** | **5** | **36606457** | **36688436** | **+** | 420 |
| SLC1A4 | 6509 | 2 | 65215579 | 65250999 | + | 145 |
| **SLC1A6** | **6511** | **19** | **15060845** | **15121455** | **-** | 503 |
| **SLC1A7** | **6512** | **1** | **53552855** | **53608304** | **-** | 472 |
| SLC38A1 | 81539 | 12 | 46576838 | 46663208 | - | 441 |
| SUCLG2 | 8801 | 3 | 67410884 | 67705038 | - | 1963 |

^All genes in table were included in the glutamate pathway gene-set. Genes marked in bold are the genes that were included in the reduced glutamate receptors/transporters gene-set (n=32). NSNPS, number of single nucleotide polymorphisms (SNPs).^

**Table S2**. Summary table of all genes in GABA gene-set included in analyses.

| Gene name | Entrez gene ID | Chromosome | Start position | End position | strand | NSNPS |
| --- | --- | --- | --- | --- | --- | --- |
| ABAT | 18 | 16 | 8768444 | 8878432 | + | 1013 |
| ADCY1 | 107 | 7 | 45614125 | 45762715 | + | 760 |
| ADCY10 | 55811 | 1 | 167778357 | 167883608 | - | 659 |
| ADCY2 | 108 | 5 | 7396343 | 7830194 | + | 2563 |
| ADCY3 | 109 | 2 | 25042038 | 25142602 | - | 694 |
| ADCY4 | 196883 | 14 | 24787555 | 24804277 | - | 81 |
| ADCY5 | 111 | 3 | 123001143 | 123167924 | - | 858 |
| ADCY6 | 112 | 12 | 49159975 | 49182820 | - | 81 |
| ADCY7 | 113 | 16 | 50278830 | 50352046 | + | 333 |
| ADCY8 | 114 | 8 | 131792546 | 132053012 | - | 1901 |
| ADCY9 | 115 | 16 | 4012650 | 4166186 | - | 1082 |
| ALDH5A1 | 7915 | 6 | 24495197 | 24537435 | + | 297 |
| ALDH9A1 | 223 | 1 | 165631449 | 165667900 | - | 239 |
| AP1B1 | 162 | 22 | 29723669 | 29784754 | - | 255 |
| AP1G2 | 8906 | 14 | 24028777 | 24038754 | - | 14 |
| AP2A1 | 160 | 19 | 50270180 | 50310369 | + | 165 |
| AP2A2 | 161 | 11 | 925809 | 1012245 | + | 487 |
| AP2B1 | 163 | 17 | 33913918 | 34053436 | + | 746 |
| AP2M1 | 1173 | 3 | 183892634 | 183901879 | + | 53 |
| AP2S1 | 1175 | 19 | 47341423 | 47354203 | - | 35 |
| CACNA1A | 773 | 19 | 13317256 | 13617274 | - | 1465 |
| CACNA1B | 774 | 9 | 140772241 | 141019076 | + | 880 |
| CACNA1C | 775 | 12 | 2079952 | 2807115 | + | 3692 |
| CACNA1D | 776 | 3 | 53529076 | 53847179 | + | 1844 |
| CACNA1E | 777 | 1 | 181452447 | 181775920 | + | 1671 |
| CACNA1F | 778 | X | 49061523 | 49089833 | - |  |
| CACNA1G | 8913 | 17 | 48638429 | 48704835 | + | 310 |
| CACNA1H | 8912 | 16 | 1203241 | 1271772 | + | 422 |
| CACNA1I | 8911 | 22 | 39966758 | 40085740 | + | 591 |
| CACNA1S | 779 | 1 | 201008635 | 201081694 | - | 505 |
| CACNA2D1 | 781 | 7 | 81575760 | 82073031 | - | 3150 |
| CACNA2D2 | 9254 | 3 | 50400230 | 50540892 | - | 656 |
| CACNA2D3 | 55799 | 3 | 54156620 | 55108584 | + | 5930 |
| CACNA2D4 | 93589 | 12 | 1901123 | 2027870 | - | 775 |
| CACNB1 | 782 | 17 | 37329709 | 37353956 | - | 89 |
| CACNB2 | 783 | 10 | 18429373 | 18830688 | + | 2968 |
| CACNB3 | 784 | 12 | 49208215 | 49222726 | + | 46 |
| CACNB4 | 785 | 2 | 152689285 | 152955593 | - | 1246 |
| CACNG1 | 786 | 17 | 65040652 | 65052913 | + | 56 |
| CACNG2 | 10369 | 22 | 36956916 | 37098690 | - | 720 |
| CACNG3 | 10368 | 16 | 24266874 | 24373737 | + | 675 |
| CACNG4 | 27092 | 17 | 64960980 | 65029518 | + | 432 |
| CACNG5 | 27091 | 17 | 64831235 | 64881941 | + | 373 |
| CACNG6 | 59285 | 19 | 54494403 | 54515920 | + | 115 |
| CACNG7 | 59284 | 19 | 54412704 | 54447195 | + | 105 |
| CACNG8 | 59283 | 19 | 54466290 | 54493469 | + | 111 |
| CATSPER1 | 117144 | 11 | 65784223 | 65793988 | - | 45 |
| CATSPER2 | 117155 | 15 | 43922772 | 43941039 | - | 63 |
| CATSPER3 | 347732 | 5 | 134303596 | 134347397 | + | 207 |
| CATSPER4 | 378807 | 1 | 26517119 | 26529033 | + | 107 |
| DNM1 | 1759 | 9 | 130965634 | 131017528 | + | 223 |
| **GABARAP** | **11337** | **17** | **7143738** | **7145753** | **-** | 5 |
| **GABBR1** | **2550** | **6** | **29570005** | **29600962** | **-** | 219 |
| **GABBR2** | **9568** | **9** | **101050364** | **101471479** | **-** | 2637 |
| **GABRA1** | **2554** | **5** | **161274197** | **161326965** | **+** | 283 |
| **GABRA2** | **2555** | **4** | **46246470** | **46392056** | **-** | 727 |
| **GABRA3** | **2556** | **X** | **151334706** | **151619831** | **-** |  |
| **GABRA4** | **2557** | **4** | **46920917** | **46996424** | **-** | 406 |
| **GABRA5** | **2558** | **15** | **27111866** | **27194357** | **+** | 158 |
| **GABRA6** | **2559** | **5** | **161112658** | **161129598** | **+** | 81 |
| **GABRB1** | **2560** | **4** | **47033295** | **47432801** | **+** | 2058 |
| **GABRB2** | **2561** | **5** | **160715426** | **160975130** | **-** | 1268 |
| **GABRB3** | **2562** | **15** | **26788693** | **27018935** | **-** | 1332 |
| **GABRD** | **2563** | **1** | **1950768** | **1962192** | **+** | 10 |
| **GABRE** | **2564** | **X** | **151121596** | **151143156** | **-** |  |
| **GABRG1** | **2565** | **4** | **46037786** | **46126082** | **-** | 496 |
| **GABRG2** | **2566** | **5** | **161494648** | **161582545** | **+** | 435 |
| **GABRG3** | **2567** | **15** | **27216429** | **27778373** | **+** | 2556 |
| **GABRP** | **2568** | **5** | **170210723** | **170241051** | **+** | 193 |
| **GABRQ** | **55879** | **X** | **151806637** | **151821825** | **+** |  |
| **GABRR1** | **2569** | **6** | **89887223** | **89941007** | **-** | 344 |
| **GABRR2** | **2570** | **6** | **89966840** | **90025018** | **-** | 405 |
| **GABRR3** | **200959** | **3** | **97705527** | **97754148** | **-** | 264 |
| GAD1 | 2571 | 2 | 171673200 | 171717661 | + | 172 |
| GAD2 | 2572 | 10 | 26505236 | 26593491 | + | 579 |
| GNA11 | 2767 | 19 | 3094408 | 3121468 | + | 144 |
| GNA12 | 2768 | 7 | 2767739 | 2883963 | - | 883 |
| GNA13 | 10672 | 17 | 63005407 | 63052920 | - | 84 |
| GNA14 | 9630 | 9 | 80037995 | 80263232 | - | 1496 |
| GNA15 | 2769 | 19 | 3136191 | 3163766 | + | 201 |
| GNAI1 | 2770 | 7 | 79764140 | 79848725 | + | 383 |
| GNAI2 | 2771 | 3 | 50264120 | 50296786 | + | 114 |
| GNAI3 | 2773 | 1 | 110091186 | 110138465 | + | 181 |
| GNAL | 2774 | 18 | 11689014 | 11885684 | + | 1003 |
| GNAO1 | 2775 | 16 | 56225251 | 56391356 | + | 866 |
| GNAQ | 2776 | 9 | 80335189 | 80646219 | - | 1344 |
| GNAS | 2778 | 20 | 57414756 | 57486250 | + | 323 |
| GNAT1 | 2779 | 3 | 50229043 | 50235129 | + | 12 |
| GNAT2 | 2780 | 1 | 110145889 | 110155705 | - | 45 |
| GNAZ | 2781 | 22 | 23412669 | 23467224 | + | 256 |
| GNB1 | 2782 | 1 | 1716725 | 1822552 | - | 250 |
| GNB1L | 54584 | 22 | 19775932 | 19842462 | - | 369 |
| GNB2 | 2783 | 7 | 100271363 | 100276792 | + | 19 |
| GNB3 | 2784 | 12 | 6949375 | 6956564 | + | 34 |
| GNB4 | 59345 | 3 | 179113876 | 179169371 | - | 290 |
| GNB5 | 10681 | 15 | 52413123 | 52483565 | - | 486 |
| GNG10 | 2790 | 9 | 114423851 | 114432526 | + | 50 |
| GNG11 | 2791 | 7 | 93551016 | 93555826 | + | 32 |
| GNG12 | 55970 | 1 | 68167149 | 68299436 | - | 702 |
| GNG13 | 51764 | 16 | 848041 | 850733 | - | 33 |
| GNG2 | 54331 | 14 | 52327022 | 52436518 | + | 794 |
| GNG3 | 2785 | 11 | 62475066 | 62476678 | + | 5 |
| GNG4 | 2786 | 1 | 235710985 | 235814054 | - | 543 |
| GNG5 | 2787 | 1 | 84964006 | 84972262 | - | 37 |
| GNG7 | 2788 | 19 | 2511218 | 2702746 | - | 1041 |
| GPHN | 10243 | 14 | 66974125 | 67648525 | + | 3011 |
| GPR37 | 2861 | 7 | 124385655 | 124406079 | - | 81 |
| KCNH2 | 3757 | 7 | 150642044 | 150675402 | - | 179 |
| KCNN1 | 3780 | 19 | 18062111 | 18110133 | + | 207 |
| KCNN2 | 3781 | 5 | 113698016 | 113832197 | + | 840 |
| KCNN3 | 3782 | 1 | 154669938 | 154842754 | - | 925 |
| KCNN4 | 3783 | 19 | 44270685 | 44286269 | - | 72 |
| KCNQ2 | 3785 | 20 | 62031561 | 62103993 | - | 607 |
| KCNQ3 | 3786 | 8 | 133133105 | 133493004 | - | 2095 |
| MRAS | 22808 | 3 | 138066490 | 138124377 | + | 307 |
| NSF | 4905 | 17 | 44668035 | 44834830 | + | 108 |
| OPN1SW | 611 | 7 | 128412543 | 128415844 | - | 20 |
| RPS27A | 6233 | 2 | 55459039 | 55462989 | + | 27 |
| SLC32A1 | 140679 | 20 | 37353105 | 37358015 | + | 20 |
| **SLC6A1** | **6529** | **3** | **11034420** | **11080935** | **+** | 267 |
| **SLC6A11** | **6538** | **3** | **10857917** | **10980146** | **+** | 739 |
| **SLC6A12** | **6539** | **12** | **299243** | **323740** | **-** | 169 |
| **SLC6A13** | **6540** | **12** | **329787** | **372039** | **-** | 322 |
| UBA52 | 7311 | 19 | 18674576 | 18688270 | + | 83 |
| UBB | 7314 | 17 | 16284367 | 16286059 | + | 7 |
| UBC | 7316 | 12 | 125396192 | 125399587 | - | 23 |
| UBD | 10537 | 6 | 29523389 | 29527702 | - | 42 |
| UBQLN1 | 29979 | 9 | 86274878 | 86323168 | - | 265 |

^All genes in table were included in the GABA pathway gene-set. Genes marked in bold are the genes that were included in the reduced GABA receptors/transporters gene-set (n=26). NSNPS, number of single nucleotide polymorphisms (SNPs).^

**Table S3:** Scanner parameters across sites

| Site | Manufacturer | Model | Software version | Acquisition sequence | Slices | TR  [s] | TE  [ms] | FA  [˚] | Coverage | Thickness  [mm] | Resolution  [mm^3^] | FOV |
| --- | --- | --- | --- | --- | --- | --- | --- | --- | --- | --- | --- | --- |
| Cambridge | Siemens | Verio | Syngo MR B17 | Tfl3d1_ns | 176 | 2.3 | 2.95 | 9 | 256*256 | 1.2 | 1.1*1.1*1.2 | 270 |
| KCL | GE Medical systems | Discovery mr750 | LX MR DV23.1_V02_1317.c | SAG ADNI GO ACC SPGR | 196 | 7.31 | 3.02 | 11 |  |  |  |  |
| Mannheim | Siemens | TimTrio | Syngo MR B17 | MPRAGE ADNI | 176 | 2.3 | 2.93 | 9 |  |  |  |  |
| Nijmegen | Siemens | Skyra | Syngo MR D13 | Tfl3d1_16ns | 176 | 2.3 | 2.93 | 9 |  |  |  |  |
| Rome | GE Medical systems | Signa HDxt | 24/LX/MR HD16.0_V02_1131.a | SAG ADNI GO ACC SPGR | 172 | 5.96 | 1.76 | 11 |  |  |  |  |
| Utrecht | Philips Medical Systems | Achieva/ Ingenia CX | 3.2.3/3.2.3.1/  5.1.9/5.1.9.1 | ADNI GO 2 | 170 | 6.76 | 3.1 | 9 |  |  |  |  |

^Abbreviations: FA, flip angle; FOV, field of view; TE, echo time; TR, repetition time.^

**Table S4:** Medication information

|  | **ASD** | **NTC** |
| --- | --- | --- |
| **N** | 122 | 20 |
| Antidepressants  SSRIs  Tetracyclic (TeCA)  Tricyclic (TCA) | 34  29  2  3 | 6  5  1  0 |
| Antiepileptics | 11 | 2 |
| Antimigraine preparations | 4 | 0 |
| Antipsychotics  Aripiprazole  Clozapine  Pipamperone  Quetiapine  Risperidone | 28  6  1  2  1  18 | 1  0  0  0  0  1 |
| Anxiolytics | 2 | 1 |
| Drugs used in Addictive Disorder | 0 | 1 |
| Hypnotics & Sedatives  Hyoscine butylbromide  Melatonin  Niaprazine | 40  1  38  1 | 2  0  2  0 |
| Other Analgesics & Antipyretics  Opioids  Others | 4  1  3 | 4  0  4 |
| Psychostimulants & Other drugs used to treat ADHD  Atomoxetine  Dexamfetamine  Methylphenidate hydrochloride | 47  3  1  43 | 9  2  0  7 |

^Note. Participants may have taken up to 3 different types of medication across the listed categories during study participation.^

**Table S5:** Glutamate and GABA and SSP subscale competitive gene-set analysis results

| **Glutamate: Pathway gene-set (N=72)** | **BETA** | **P** | | **P_FDR_** | **SE** |
| --- | --- | --- | --- | --- | --- |
| SSP Auditory filtering | 0.123 | 0.111 | | 0.147 | 0.101 |
| SSP Low energy/weak | 0.123 | 0.112 | | 0.147 | 0.101 |
| SSP Movement sensitivity | 0.120 | 0.117 | | 0.147 | 0.101 |
| SSP Tactile sensitivity | 0.109 | 0.139 | | 0.147 | 0.101 |
| SSP Taste/smell sensitivity | 0.106 | 0.147 | | 0.147 | 0.101 |
| SSP Underresponsive/seeks attention | 0.109 | 0.140 | | 0.147 | 0.101 |
| SSP Visual/auditory sensitivity | 0.123 | 0.111 | | 0.147 | 0.101 |
| **Glutamate: Receptors/transporters gene-set (N=31)** |  |  | |  |  |
| SSP Auditory filtering | 0.212 | | 0.089 | 0.138 | 0.157 |
| SSP Low energy/weak | 0.212 | | 0.089 | 0.138 | 0.157 |
| SSP Movement sensitivity | 0.172 | | 0.138 | 0.138 | 0.157 |
| SSP Tactile sensitivity | 0.181 | | 0.126 | 0.138 | 0.157 |
| SSP Taste/smell sensitivity | 0.184 | | 0.121 | 0.138 | 0.158 |
| SSP Underresponsive/seeks attention | 0.181 | | 0.125 | 0.138 | 0.157 |
| SSP Visual/auditory sensitivity | 0.213 | | 0.088 | 0.138 | 0.157 |
| **GABA: Pathway gene-set (N=124)** | **BETA** | | **P** | **PFDR** | **SE** |
| **SSP Auditory filtering** | **0.157** | | **0.023** | **0.024** | **0.079** |
| **SSP Low energy/weak** | **0.157** | | **0.023** | **0.024** | **0.079** |
| **SSP Movement sensitivity** | **0.166** | | **0.018** | **0.024** | **0.079** |
| **SSP Tactile sensitivity** | **0.157** | | **0.024** | **0.024** | **0.079** |
| **SSP Taste/smell sensitivity** | **0.157** | | **0.024** | **0.024** | **0.079** |
| **SSP Underresponsive/seeks attention** | **0.157** | | **0.023** | **0.024** | **0.079** |
| **SSP Visual/auditory sensitivity** | **0.157** | | **0.023** | **0.024** | **0.079** |
| **GABA: Receptors/transporters gene-set (N=23)** |  | |  |  |  |
| **SSP Auditory filtering** | **0.382** | | **0.027** | **0.027** | **0.198** |
| **SSP Low energy/weak** | **0.382** | | **0.027** | **0.027** | **0.198** |
| **SSP Movement sensitivity** | **0.453** | | **0.011** | **0.027** | **0.198** |
| **SSP Tactile sensitivity** | **0.417** | | **0.018** | **0.027** | **0.198** |
| **SSP Taste/smell sensitivity** | **0.405** | | **0.021** | **0.027** | **0.199** |
| **SSP Underresponsive/seeks attention** | **0.416** | | **0.018** | **0.027** | **0.198** |
| **SSP Visual/auditory sensitivity** | **0.383** | | **0.027** | **0.027** | **0.198** |

^N, number of genes in analysis. Diagnosis was indicated as a binary variable. SSP, Short Sensory Profile; P^_FDR_ ^p-value corrected using False discovery rate (FDR); SE, standard error of the regression coefficient. Significant results (p^_FDR_^<0.05) marked in bold.^

**Competitive gene-set analysis on cortical thickness**

**Table S6**. Glutamate pathway - left hemisphere competitive gene-set analysis

| FreeSurfer region | NGENES | BETA | *P* | *P*_FDR_ |
| --- | --- | --- | --- | --- |
| Banks of superior temporal sulcus | 72 | -0.028 | 0.603 | 0.967 |
| **Caudal anterior cingulate cortex** | **72** | **0.182** | **0.042** | 0.776 |
| Caudal middle frontal gyrus | 72 | -0.047 | 0.671 | 0.967 |
| Cuneus | 72 | -0.090 | 0.809 | 0.967 |
| Entorhinal cortex | 72 | 0.010 | 0.461 | 0.967 |
| Frontal pole | 72 | 0.030 | 0.388 | 0.967 |
| Fusiform gyrus | 72 | -0.045 | 0.669 | 0.967 |
| Inferior parietal cortex | 72 | -0.090 | 0.801 | 0.967 |
| Inferior temporal gyrus | 72 | -0.115 | 0.867 | 0.967 |
| Insula | 72 | 0.030 | 0.387 | 0.967 |
| Isthmus-cingulate cortex | 72 | -0.125 | 0.883 | 0.967 |
| Lateral occipital gyrus | 72 | -0.107 | 0.851 | 0.967 |
| Lateral orbital frontal cortex | 72 | 0.164 | 0.064 | 0.776 |
| Lingual gyrus | 72 | 0.105 | 0.157 | 0.967 |
| Medial orbital frontal cortex | 72 | 0.075 | 0.244 | 0.967 |
| Middle temporal gyrus | 72 | -0.056 | 0.698 | 0.967 |
| Paracentral lobule | 72 | 0.046 | 0.330 | 0.967 |
| Parahippocampal gyrus | 72 | -0.030 | 0.611 | 0.967 |
| Pars opercularis | 72 | -0.077 | 0.765 | 0.967 |
| Pars orbitalis | 72 | 0.157 | 0.068 | 0.776 |
| Pars triangularis | 72 | -0.054 | 0.694 | 0.967 |
| Pericalcarine cortex | 72 | -0.001 | 0.503 | 0.967 |
| Postcentral gyrus | 72 | -0.200 | 0.971 | 0.971 |
| Posterior cingulate cortex | 72 | -0.088 | 0.792 | 0.967 |
| Precentral gyrus | 72 | -0.045 | 0.664 | 0.967 |
| Precuneus cortex | 72 | -0.012 | 0.548 | 0.967 |
| Rostral anterior cingulate cortex | 72 | -0.130 | 0.888 | 0.967 |
| Rostral middle frontal gyrus | 72 | -0.038 | 0.640 | 0.967 |
| Superior frontal gyrus | 72 | -0.057 | 0.702 | 0.967 |
| Superior parietal cortex | 72 | -0.051 | 0.686 | 0.967 |
| Superior temporal gyrus | 72 | -0.160 | 0.936 | 0.967 |
| Supramarginal gyrus | 72 | -0.166 | 0.939 | 0.967 |
| Temporal pole | 72 | -0.047 | 0.667 | 0.967 |
| Transverse temporal cortex | 72 | -0.106 | 0.846 | 0.967 |

^NGENES, number of genes in analysis. The associations with significance are marked in bold.^

**Table S7**. Glutamate receptors/transporters - left hemisphere competitive gene-set analysis

| FreeSurfer region | NGENES | BETA | P | P_FDR_ |
| --- | --- | --- | --- | --- |
| Banks of superior temporal sulcus | 31 | -0.104 | 0.737 | 0.983 |
| Caudal anterior cingulate cortex | 31 | 0.043 | 0.398 | 0.983 |
| Caudal middle frontal gyrus | 31 | 0.099 | 0.273 | 0.983 |
| Cuneus | 31 | -0.120 | 0.771 | 0.983 |
| Entorhinal cortex | 31 | 0.083 | 0.307 | 0.983 |
| Frontal pole | 31 | -0.067 | 0.657 | 0.983 |
| Fusiform gyrus | 31 | -0.133 | 0.796 | 0.983 |
| Inferior parietal cortex | 31 | -0.098 | 0.724 | 0.983 |
| Inferior temporal gyrus | 31 | -0.309 | 0.973 | 0.983 |
| Insula | 31 | -0.113 | 0.759 | 0.983 |
| Isthmus-cingulate cortex | 31 | 0.045 | 0.391 | 0.983 |
| Lateral occipital gyrus | 31 | -0.178 | 0.865 | 0.983 |
| Lateral orbital frontal cortex | 31 | -0.007 | 0.516 | 0.983 |
| Lingual gyrus | 31 | -0.125 | 0.777 | 0.983 |
| Medial orbital frontal cortex | 31 | -0.221 | 0.904 | 0.983 |
| Middle temporal gyrus | 31 | -0.027 | 0.563 | 0.983 |
| **Paracentral lobule** | **31** | **0.335** | **0.020** | 0.676 |
| Parahippocampal gyrus | 31 | -0.048 | 0.615 | 0.983 |
| Pars opercularis | 31 | -0.302 | 0.966 | 0.983 |
| Pars orbitalis | 31 | 0.074 | 0.326 | 0.983 |
| Pars triangularis | 31 | -0.330 | 0.976 | 0.983 |
| Pericalcarine cortex | 31 | 0.147 | 0.184 | 0.983 |
| Postcentral gyrus | 31 | -0.119 | 0.767 | 0.983 |
| Posterior cingulate cortex | 31 | -0.068 | 0.657 | 0.983 |
| Precentral gyrus | 31 | 0.001 | 0.497 | 0.983 |
| Precuneus cortex | 31 | -0.188 | 0.877 | 0.983 |
| Rostral anterior cingulate cortex | 31 | -0.355 | 0.983 | 0.983 |
| Rostral middle frontal gyrus | 31 | -0.268 | 0.947 | 0.983 |
| Superior frontal gyrus | 31 | -0.253 | 0.934 | 0.983 |
| Superior parietal cortex | 31 | -0.270 | 0.952 | 0.983 |
| Superior temporal gyrus | 31 | -0.171 | 0.852 | 0.983 |
| Supramarginal gyrus | 31 | -0.093 | 0.711 | 0.983 |
| Temporal pole | 31 | -0.039 | 0.592 | 0.983 |
| Transverse temporal cortex | 31 | -0.011 | 0.528 | 0.983 |

^NGENES, number of genes in analysis. The associations with significance are marked in bold.^

**Table S8**. GABA pathway - left hemisphere competitive gene-set analysis

| FreeSurfer region | NGENES | BETA | P | P_FDR_ |
| --- | --- | --- | --- | --- |
| Caudal anterior cingulate cortex | 124 | -0.021 | 0.599 | 0.956 |
| Caudal middle frontal gyrus | 124 | 0.086 | 0.150 | 0.555 |
| Cuneus | 124 | 0.066 | 0.211 | 0.652 |
| Entorhinal cortex | 124 | -0.090 | 0.867 | 0.956 |
| Frontal pole | 124 | 0.084 | 0.154 | 0.555 |
| Fusiform gyrus | 124 | 0.119 | 0.074 | 0.458 |
| Inferior parietal cortex | 124 | -0.046 | 0.716 | 0.956 |
| Inferior temporal gyrus | 124 | -0.079 | 0.829 | 0.956 |
| Insula | 124 | 0.017 | 0.415 | 0.940 |
| Isthmus-cingulate cortex | 124 | -0.101 | 0.894 | 0.956 |
| Lateral occipital gyrus | 124 | 0.036 | 0.331 | 0.812 |
| Lateral orbital frontal cortex | 124 | -0.070 | 0.806 | 0.956 |
| Lingual gyrus | 124 | 0.082 | 0.163 | 0.555 |
| Medial orbital frontal cortex | 124 | -0.078 | 0.829 | 0.956 |
| **Middle temporal gyrus** | **124** | **0.227** | **0.004** | 0.127 |
| Paracentral lobule | 124 | -0.145 | 0.956 | 0.956 |
| Parahippocampal gyrus | 124 | -0.004 | 0.519 | 0.956 |
| Pars opercularis | 124 | -0.047 | 0.715 | 0.956 |
| Pars orbitalis | 124 | 0.094 | 0.129 | 0.555 |
| Pars triangularis | 124 | 0.116 | 0.081 | 0.458 |
| **Pericalcarine cortex** | **124** | **0.172** | **0.020** | 0.347 |
| Postcentral gyrus | 124 | -0.073 | 0.813 | 0.956 |
| Posterior cingulate cortex | 124 | -0.057 | 0.755 | 0.956 |
| Precentral gyrus | 124 | 0.004 | 0.482 | 0.956 |
| Precuneus cortex | 124 | -0.078 | 0.825 | 0.956 |
| Rostral anterior cingulate cortex | 124 | -0.030 | 0.643 | 0.956 |
| Rostral middle frontal gyrus | 124 | 0.118 | 0.080 | 0.458 |
| **Superior frontal gyrus** | **124** | **0.155** | **0.031** | 0.352 |
| Superior parietal cortex | 124 | 0.036 | 0.335 | 0.812 |
| Superior temporal gyrus | 124 | -0.054 | 0.746 | 0.956 |
| Caudal anterior cingulate cortex | 124 | -0.114 | 0.918 | 0.956 |
| Supramarginal gyrus | 124 | -0.129 | 0.937 | 0.956 |
| Temporal pole | 124 | 0.049 | 0.280 | 0.793 |
| Transverse temporal cortex | 124 | -0.094 | 0.875 | 0.956 |

^NGENES, number of genes in analysis. The associations with significance are marked in bold.^

**Table S9**. GABA receptors/transporters - left hemisphere competitive gene-set analysis

| FreeSurfer region | NGENES | BETA | P | P_FDR_ |
| --- | --- | --- | --- | --- |
| Caudal anterior cingulate cortex | 23 | 0.005 | 0.496 | 0.849 |
| Caudal middle frontal gyrus | 23 | -0.123 | 0.724 | 0.849 |
| Cuneus | 23 | 0.133 | 0.260 | 0.849 |
| Entorhinal cortex | 23 | 0.032 | 0.437 | 0.849 |
| Frontal pole | 23 | -0.113 | 0.707 | 0.849 |
| Fusiform gyrus | 23 | 0.166 | 0.211 | 0.849 |
| Inferior parietal cortex | 23 | -0.034 | 0.568 | 0.849 |
| Inferior temporal gyrus | 23 | -0.175 | 0.800 | 0.907 |
| Insula | 23 | 0.092 | 0.325 | 0.849 |
| Isthmus-cingulate cortex | 23 | 0.013 | 0.475 | 0.849 |
| Lateral occipital gyrus | 23 | 0.138 | 0.251 | 0.849 |
| Lateral orbital frontal cortex | 23 | -0.244 | 0.886 | 0.921 |
| Lingual gyrus | 23 | -0.253 | 0.885 | 0.921 |
| Medial orbital frontal cortex | 23 | -0.018 | 0.535 | 0.849 |
| **Middle temporal gyrus** | **23** | **0.535** | **0.006** | 0.205 |
| Paracentral lobule | 23 | -0.117 | 0.708 | 0.849 |
| Parahippocampal gyrus | 23 | 0.037 | 0.429 | 0.849 |
| Pars opercularis | 23 | 0.022 | 0.457 | 0.849 |
| Pars orbitalis | 23 | 0.308 | 0.069 | 0.647 |
| Pars triangularis | 23 | -0.084 | 0.657 | 0.849 |
| Pericalcarine cortex | 23 | 0.172 | 0.208 | 0.849 |
| Postcentral gyrus | 23 | -0.064 | 0.621 | 0.849 |
| Posterior cingulate cortex | 23 | 0.074 | 0.359 | 0.849 |
| Precentral gyrus | 23 | 0.292 | 0.084 | 0.647 |
| Precuneus cortex | 23 | -0.101 | 0.686 | 0.849 |
| Rostral anterior cingulate cortex | 23 | 0.266 | 0.095 | 0.647 |
| Rostral middle frontal gyrus | 23 | 0.382 | 0.034 | 0.586 |
| Superior frontal gyrus | 23 | -0.102 | 0.687 | 0.849 |
| Superior parietal cortex | 23 | 0.106 | 0.307 | 0.849 |
| Superior temporal gyrus | 23 | -0.256 | 0.894 | 0.921 |
| Caudal anterior cingulate cortex | 23 | 0.013 | 0.476 | 0.849 |
| Supramarginal gyrus | 23 | -0.117 | 0.711 | 0.849 |
| Temporal pole | 23 | -0.094 | 0.670 | 0.849 |
| Transverse temporal cortex | 23 | -0.307 | 0.933 | 0.933 |

^NGENES, number of genes in analysis. The associations with significance are marked in bold.^

**Expression profiles**

**Separate age-groups.** LEAP sample separated into children (6-11 years) adolescents (12-17 years) and adults (18 years and older). As we know that cortical thickness (CT) is strongly associated with age (1), and we found the interregional profiles of group differences in CT (autism-NTC) to be associated with the interregional profile of gene expression for both glutamate and GABA-pathway gene-sets, we wanted to investigate whether these correlations differ across age groups. In children (autism = 52, NTC = 64), there were no significant effects (glu-pathway: t=0.46, p_FDR_ = 0.93, d=0.17, GABA-pathway: t=-0.09, p_FDR_ = 0.93, d=-0.03), as can be seen in Figure S1. In adolescents (autism = 92, NTC = 88), there were significant effects in the opposite direction of the findings in the whole sample (glu-pathway: t=3.72, p_FDR_ = 0.002, d=1.41, GABA-pathway: t=2.78, p_FDR_ = 0.01, d=0.87), as can be seen in Figure S2. These results suggest that decreased group differences in interregional profiles of CT in adolescents are associated with the interregional profiles of gene expression in both glutamate and GABA-pathway gene sets. In adults (autism = 135, NTC = 102), the significant associations were similar to the whole sample (glu-pathway: t=-2.49, p_FDR_ = 0.04, d=-0.94, GABA-pathway: t=-2.10, p_FDR_ = 0.04, d=-0.66), where regions with greater gene expression of glutamate and GABAergic pathway genes show greater differences in CT. These results suggests that the effect found across the whole sample are driven by adults.

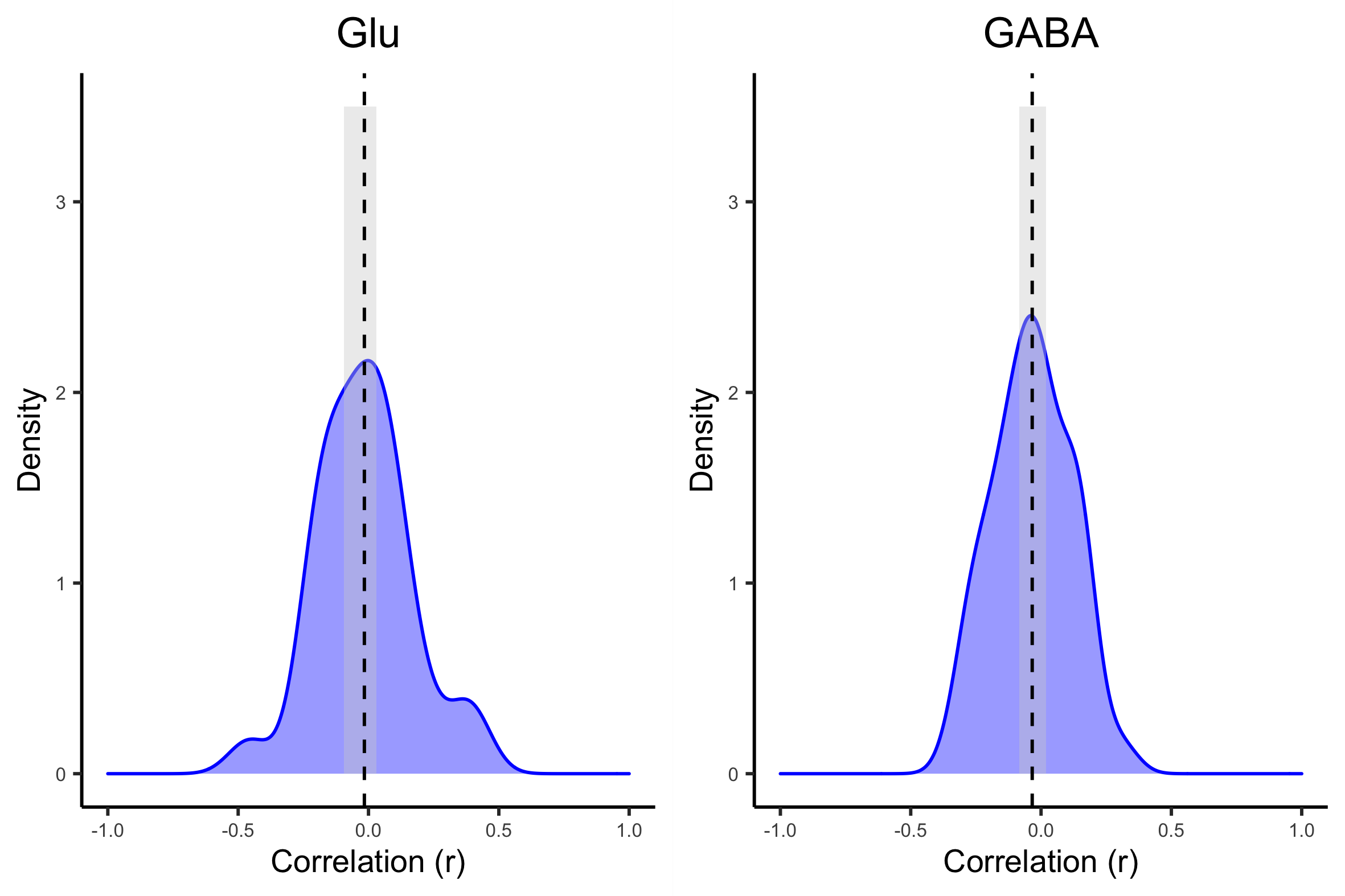

**Figure S1,** **Children:** Distributions of the inter-regional correlation coefficients between differences in cortical thickness (CT) and profiles of gene-expression. The CT-difference profile was obtained from children aged 6-11 in our LEAP data, and the expression profiles from the Allen Human Brian Atlas (AHBA), in our glutamate-pathway and GABA-pathway gene-sets. The x-axes show the correlation coefficient between CT-difference and expression profile for the gene-set; the y-axes show the estimated probability density for the correlation coefficients; the vertical dashed-lines indicate the average expression-CT difference correlation coefficient across all the marker genes in a gene-set; and the edges of the gray boxes indicate the 2.5% and 97.5%-critical values obtained from the empirical null distribution of the average expression-thickness correlation coefficient. If a vertical line sits outside the gray box, it implies that there is a significant association between gene-set and CT at the unadjusted 5% significance level.

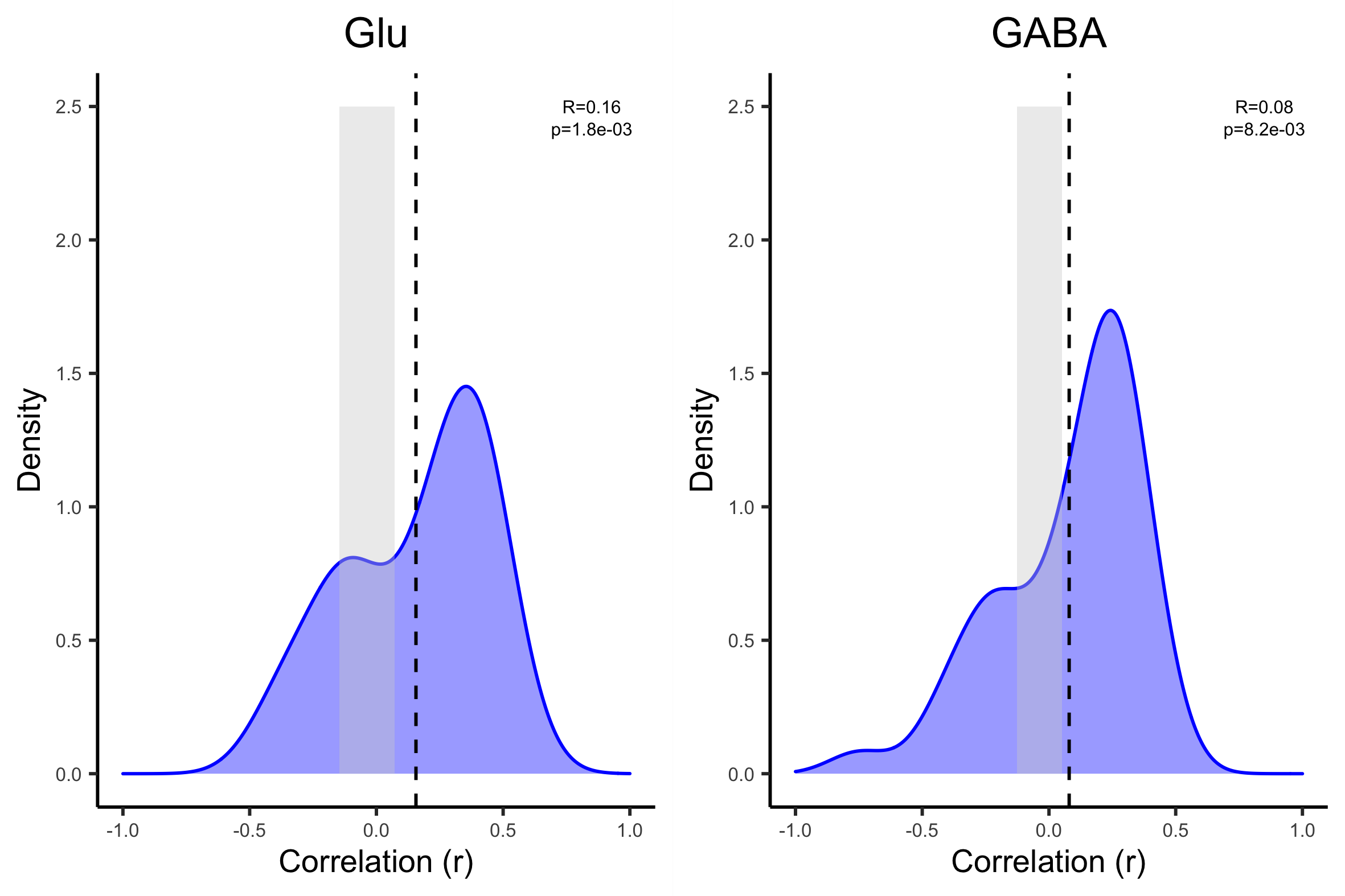

**Figure S2,** **Adolescents:** Distributions of the inter-regional correlation coefficients between differences in cortical thickness (CT) and profiles of gene-expression. The CT-difference profile was obtained from adolescents aged 12-18 in our LEAP data, and the expression profiles from the Allen Human Brian Atlas (AHBA), in our glutamate-pathway and GABA-pathway gene-sets. The x-axes show the correlation coefficient between CT-difference and expression profile for the gene-set; the y-axes show the estimated probability density for the correlation coefficients; the vertical dashed-lines indicate the average expression-CT difference correlation coefficient across all the marker genes in a gene-set; and the edges of the gray boxes indicate the 2.5% and 97.5%-critical values obtained from the empirical null distribution of the average expression-thickness correlation coefficient. If a vertical line sits outside the gray box, it implies that there is a significant association between gene-set and CT at the unadjusted 5% significance level.

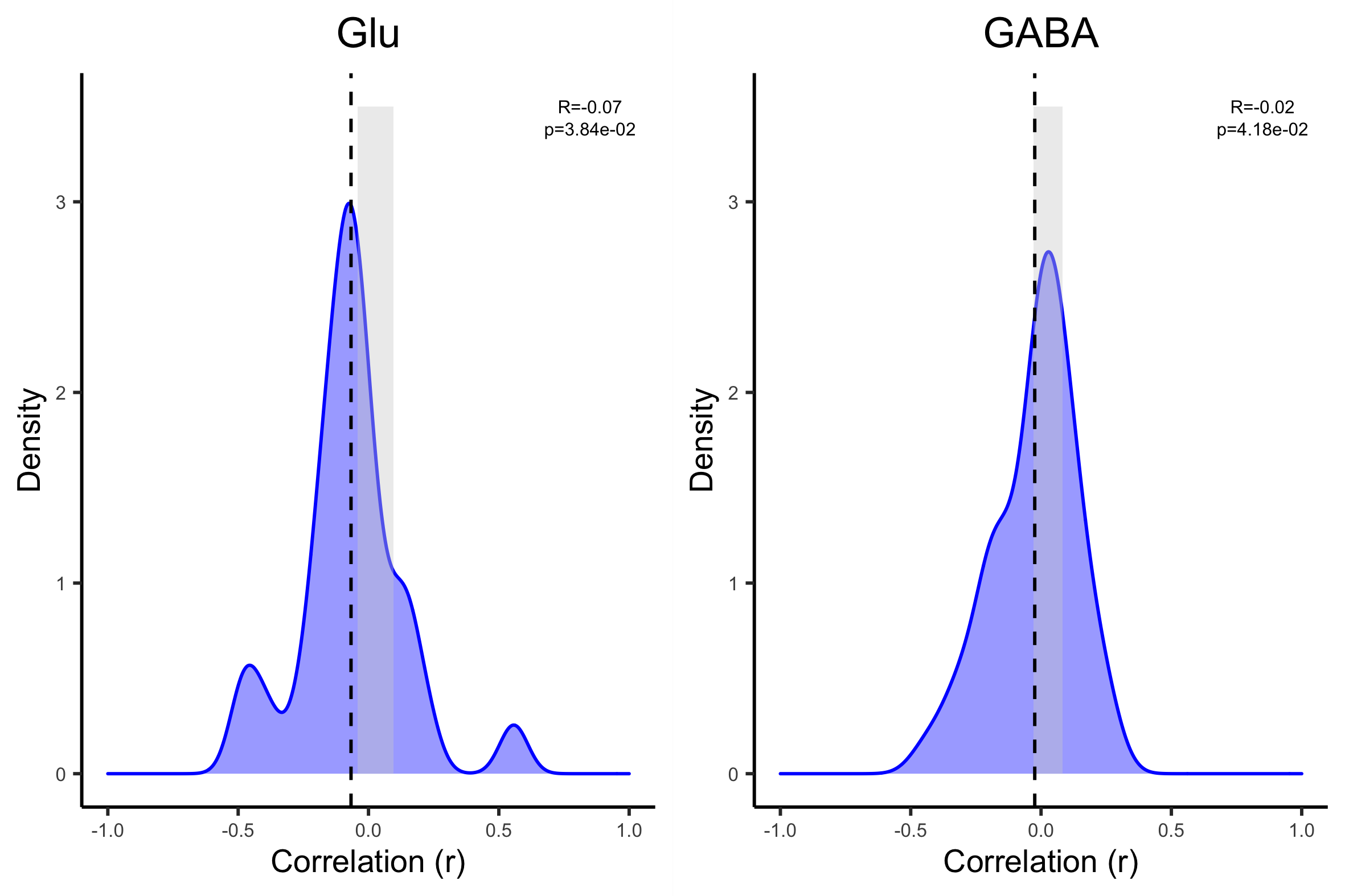

**Figure S3, Adults**: Distributions of the inter-regional correlation coefficients between differences in cortical thickness (CT) and profiles of gene-expression. The CT-difference profile was obtained from adults aged 18-30 in our LEAP data, and the expression profiles from the Allen Human Brian Atlas (AHBA), in our glutamate-pathway and GABA-pathway gene-sets. The x-axes show the correlation coefficient between CT-difference and expression profile for the gene-set; the y-axes show the estimated probability density for the correlation coefficients; the vertical dashed-lines indicate the average expression-CT difference correlation coefficient across all the marker genes in a gene-set; and the edges of the gray boxes indicate the 2.5% and 97.5%-critical values obtained from the empirical null distribution of the average expression-thickness correlation coefficient. If a vertical line sits outside the gray box, it implies that there is a significant association between gene-set and CT at the unadjusted 5% significance level.

**Replicating gene-expression analysis using ABIDE data.** The ABIDE cortical thickness (CT) data was acquired from the open source data base (<http://fcon_1000.projects.nitrc.org/indi/abide/>) (2), where we selected participants within the same age-range as our LEAP sample (6-30 years, matched for age, sex and IQ). To replicate the analysis performed in our LEAP sample, we calculated CT-difference scores between the autism and neurotypical control group (NTC) as described in the manuscript.

Gene expression analysis using the ABIDE CT-difference interregional profiles showed no significant associations between glutamate (t=0.21, p =084, d=0.07) and GABA (t=-0.63, p = 0.84, d=0.20) gene expressions and cortical thickness. As seen in Figure S3, the correlation coefficients are not significant, however the GABA results points towards a similar effect as seen in the LEAP sample (seen in Figure 1 in the manuscript). Figure S4 shows the variations in CT and the most significant gene in each gene set (the gene with the most negative correlation coefficient), illustrating the similarity between the inter-regional variations in CT and the expression of the gene.

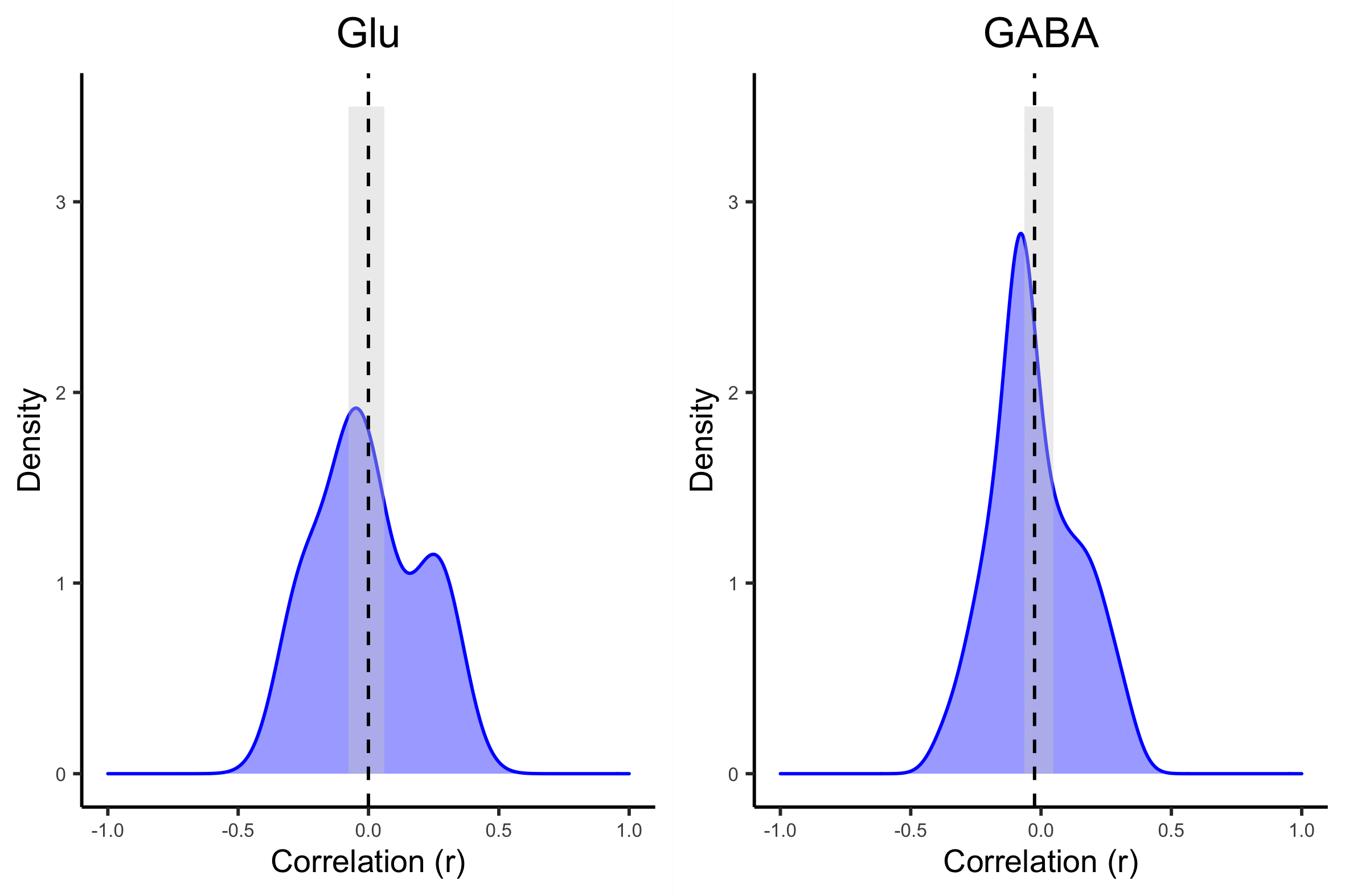

**FIGURE S4.** Distributions of the inter-regional correlation coefficients between differences in cortical thickness (CT) and profiles of gene-expression. The CT-difference profile was obtained from the ABIDE data, and the expression profiles from the Allen Human Brian Atlas (AHBA), in our glutamate-pathway and GABA-pathway gene-sets. The x-axis show the correlation coefficient between CT-difference and expression profile for the gene-set; the y-axis show the estimated probability density for the correlation coefficients; the vertical dashed-line indicates the average expression-CT difference correlation coefficient across all the marker genes in a gene-set; and the edges of the gray box indicates the 2.5% and 97.5%-critical values obtained from the empirical null distribution of the average expression-thickness correlation coefficient. If a vertical line sits outside the gray box, it implies that there is a significant association between gene-set and CT at the unadjusted 5% significance level.
